# Fusion Pore Formation Observed During SNARE-Mediated Vesicle Fusion with Pore-Spanning Membranes

**DOI:** 10.1101/2020.01.16.909044

**Authors:** P. Mühlenbrock, K. Herwig, L. Vuong, I. Mey, C. Steinem

## Abstract

Planar pore-spanning membranes (PSMs) have been shown to be a versatile tool to resolve docking and elementary steps of the fusion process with single large unilamellar vesicles (LUVs). However, in previous studies, we monitored only lipid mixing and did not gather information about the formation of fusion pores. To address this important step of the fusion process, we entrapped sulforhodamine B at self-quenching concentrations into LUVs containing the v-SNARE synaptobrevin 2, which were docked and fused with lipid-labeled PSMs containing the t-SNARE acceptor complex ΔN49 prepared on porous silicon substrates. By dual color spinning disc fluorescence microcopy with a time resolution of 20 ms, we could unambiguously distinguish between bursting vesicles and fusion pore formation. Owing to the aqueous compartment underneath the PSMs, vesicle bursting turned out to be an extremely rare event (< 0.01 %). From the time-resolved dual color fluorescence time traces, we were able to identify different fusion pathways including remaining three-dimensional postfusion structures with released content and flickering fusion pores. Our results on fusion pore formation and lipid diffusion from the PSM into the fusing vesicle let us conclude that the content release, i.e., fusion pore formation follows the merger of the two lipid membranes by only about 40 ms.

**STATEMENT OF SIGNIFICANCE:** Despite great efforts to develop *in vitro* fusion assays to understand the process of neuronal fusion, there is still a huge demand to provide single vesicle fusion assays that simultaneously report on all intermediate states including three-dimensional postfusion structures and fusion pore formation including flickering pores without the underlying artifact of vesicle bursting. Here, we show that pore-spanning membranes (PSMs) are ideal candidates to fulfill these demands. Owing to their planarity and the second aqueous compartments, they are readily accessible by fluorescence microscopy and provide sufficient space so that vesicle bursting becomes negligible. Dual color fluorescence microscopy allows distinguishing between different fusion intermediates and fusion pathways such as “kiss and run” fusion as well as flickering fusion pores.

## INTRODUCTION

SNARE-mediated membrane fusion has been established as one of the key steps in neuronal signal transmission and SNAREs are the engine of the fusion machinery (1–3). The highly exergonic nature of the assembly of the SNARE complex provides the energy for overcoming the barrier or even several barriers dependent on the intermediate states barrier (4). The critical step in the process of neurotransmitter release into the presynaptic cleft via exocytosis is the creation of a fusion pore that allows the content to be released (5). Exocytic fusion pores have been shown to be highly dynamic and determine the amount, size, and the kinetics of cargo release, with important consequences for downstream events (6–8). To elucidate the process of neuronal fusion with a focus on fusion pore formation, *in vitro* systems are highly desirable. Several *in vitro* assays have been developed over the past two decades (9,10). Bulk fusion assays based on lipid mixing (11,12) or content mixing (13) are straightforward to measure fusion efficiencies as well as kinetics (14) and in particular cases even intermediate states (15). Observation of SNARE-driven fusion at the single vesicle level however, has the potential to dissect and characterize fusion intermediates as well as their lifetime and kinetics much more directly. Thus, single vesicle assays have been realized based on two vesicle populations with one population being immobilized on a support (16). Fusion was defined by the fluorescent readout of lipid mixing. Besides the fact that this setup does not reflect the membrane geometry found in nature, where a highly curved vesicle fuses with a planar presynaptic membrane, it was shown that if this assay type is expanded to content mixing (fusion pore formation), a significantly reduced full fusion efficiency was observed (17,18). To account for the planar membrane geometry of the presynaptic membrane, SNARE-containing supported lipid bilayers (SLBs) (19–23) appear to be an alternative onto which individual vesicles can fuse. However, as there is close proximity between the membrane and the support resulting in a strong interaction, protocols of SLB formation need to be carefully adapted to ensure that the membrane components are still sufficiently mobile (24,25). Moreover, some of the assays based on SLBs suffer from the fact that the fusion process is SNAP25 independent (23,26). To monitor fusion pore formation during the fusion of a single vesicle with an SLB, water-soluble dyes are entrapped into the v-SNARE containing vesicles. Owing to the thin aqueous space between the bilayer and the support, the water-soluble dye gets preferentially released into the space above the bilayer (vesicle bursting), rather than into the thin aqueous space (21). To increase the distance between membrane and support, polymer cushions were implemented into the system, which still do not provide a large aqueous phase between membrane and support (20,27,28).

An alternative approach, where second aqueous compartments are provided that can take up the vesicle’s content are pore-spanning membranes (PSMs) (29,30). In previous studies, we and others have shown that this model membrane is suited to investigate SNARE mediated fusion on the single vesicle level (29,31–33). Based on the fluorescent readout, we were able to observe the diffusion of docked vesicles as well as fusion intermediates and postfusion structures in a time-resolved manner (29,31,32).

Here, we asked the question how and when fusion pore formation occurs during the elementary steps of the fusion process that can be readily identified in this system by a lipid dye with high time-resolution (29). To address this question, we generated fluorescently labeled PSMs containing the t-SNARE acceptor complex ΔN49 on porous substrates, to which synaptobrevin 2 doped large unilamellar vesicles entrapping a water-soluble dye at self-quenching concentrations were added. Dual color spinning disc fluorescence microcopy was used to monitor docking and fusion of individual vesicles with high temporal resolution (~20 ms). The porous mesh allowed us to directly observe the dye transfer into the underlying aqueous compartment. Simultaneously, we were able to measure lipid mixing enabling us to visualize different fusion pathways based on the interplay of fusion pore formation and the lipid diffusion from the PSM into the fusing vesicle.

## MATERIALS AND METHODS

### Materials

Porous SiO_2_/Si_3_N_4_ substrates with a pore diameter of 1.2 and 5.0 μm were purchased from Aquamarijn (Zutphen, the Netherlands). 1,2-dioleoyl-*sn*-glycero-3-phosphocholine (DOPC), 1-pal-mitoyl-2-oleoyl-*sn*-glycero-3-phospho-l-serine (POPS) and 1-palmitoyl-2-oleoyl-*sn*-glycero-3-phosphoethanolamine (POPE) were from Avanti Polar Lipids (Alabaster, AL, USA). Cholesterol, sulforhodamine B acid chloride (SRB) and 6-mer-capto-1-hexanol were purchased from Sigma-Aldrich (Taufkirchen, Germany). Atto488 maleimide and Atto655/488/390-1,2-dipalmitoyl-*sn*-glycero-3-phosphoethanolamine (Atto655/488/390 DPPE) were from Atto-Tec (Siegen, Germany).

### Protein expression, isolation and labeling

Expression and purification of the SNAREs was performed according to a protocol described previously (15,34). Briefly, recombinant expression of the t-SNAREs syntaxin 1A (amino acids (aa) 183-288), SNAP25a (aa 1-106 with all cysteine residues replaced by serine), the synaptobrevin 2 fragments (aa 1-96, aa 49-96, aa 49-96 S79C) and full-length synaptobrevin 2 (aa 1-116) originating from *Rattus norvegicus* was performed using transformed *Escherichia coli* BL21(DE3) cells carrying a pET28a vector. As all proteins are equipped with a His_6_-tag, purification was achieved using Ni^2+^-NTA agarose affinity chromatography, thrombin cleavage of the His6-tag overnight and final ion exchange chromatography with a MonoQ or MonoS column (Äkta purifying system, GE Healthcare, Little Chalfont, United Kingdom). Synaptobrevin 2 49-96 S79C was labeled with Atto488 maleimide. Acceptor complex ΔN49 was assembled from syntaxin 1A, SNAP25a and synaptobrevin 2 (aa 49-96, unlabeled or Atto488-labeled) and purified by ion exchange chromatography on a MonoQ column as described previously (29,31,32).

### Protein reconstitution

A co-micellization procedure in presence of *n*-oc-tyl-β-d-glycoside (*n*-OG) followed by detergent removal via size exclusion chromatography was used to reconstitute the proteins into small unilamellar vesicles (SUVs) as described previously (35). Briefly, lipids (DOPC/POPE/POPS/cholesterol; 5/2/1/2 (*n*/*n*), 0.465 mg total) were mixed in chloroform, the solvent was removed under a nitrogen stream at 30 °C and the resulting lipid film subsequently dried *in vacuo* for 2 h at room temperature. Lipid films were rehydrated with 50 μL buffer A (20 mM HEPES, 100 mM KCL, 1 mM dithiothreitol, 217 mOsm, pH 7.4) and *n*-OG for 30 min at 0 °C. Mixing with protein solution results in a final protein to lipid ratio (p/l) of 1:500 at 75 mM *n*-OG. After incubation for 45 min at 0 °C, detergent was removed by size exclusion chromatography (illustra NAP-10 G25 column; GE Healthcare) in buffer A. A second size exclusion chromatography step was performed in ultrapure water to remove remaining detergent and salt. For the preparation of giant unilamellar vesicles (GUVs) containing the ΔN49 complex, proteo-SUVs were dried on indium tin oxide slides. GUVs were produced via electroformation (3 h, 1.6 V_peak to peak_, 12 Hz) in 255 mOsm sucrose solution. For large unilamellar vesicles (LUVs) containing synaptobrevin 2, proteo-SUVs were dried in a round bottom flask in a desiccator over a saturated NaCl solution at 4 °C. Sulforhodamine B (SRB) was dissolved in buffer A (43 mM, 255 mOsm), the pH was adjusted to pH 7.4 and added to the proteo-lipid film. After incubation for 30 min, the suspended lipid film was extruded 31 times through a 400 nm polycarbonate membrane using a miniextruder (LiposoFast-Basic, Avestin, Ottawa, Ontario, Canada) resulting in a mean vesicle size of 240 ± 100 nm and quantitative protein reconstitution as determined previously (29). Stable inclusion of SRB into the vesicle lumen was verified by UV-Vis spectroscopy (Fig. S1A). The fusogenicity of the SRB containing LUVs with syn-aptobrevin 2 and SUVs containing the ΔN49 complex was analyzed using a bulk fusion assay (Fig. S1B).

### Reconstitution efficiency of ΔN49 complex in GUVs

The reconstitution efficiency *R* of the ΔN49 complex was determined according to Aimon et al. (36) with slight modifications. *R* is defined as (eq. 1):

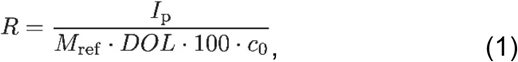

with *I*_p_, the peak membrane fluorescence of ΔN49-Atto488 doped GUVs in detector counts and *c*_0_ the nominal protein concentration of 0.1 mol%. *M*_ref_ = 47300 counts/mol% is the slope of a calibration curve of peak membrane intensities of five known Atto488 DPPE concentrations measured at exactly the same experimental conditions (Fig. S2A), whereas *DOL* is the degree of labeling of the protein determined to be 35 %. The fusion activity of the labeled ΔN49 complex was analyzed in a bulk fusion assay (Fig. S2B).

### PSM preparation

Porous SiO_2_/Si_3_N_4_ substrates with a pore diameter of 1.2 μm for single vesicle fusion experiments and 5 μm for fluorescence recovery after photobleaching (FRAP) experiments were flushed with a nitrogen stream and cleaned with a combination of argon and oxygen plasma. The surface was coated with titanium (20 s, 40 mA, 0.4 mbar, Cressington sputter coater 108 auto, Elektronen-Optik-Service, Dortmund, Germany) and subsequently with a 40 nm thick gold layer by thermal evaporation (Bal-Tec Med 020, Balzers, Liechtenstein). The substrates were functionalized in 1 mM *n*-propanolic 6-mercapto-1-hexanol solution overnight at 8 °C. For PSM preparation, substrates were rinsed with ethanol, buffer B (20 mM HEPES, 121 mM KCl, 1 mM dithiothreitol, 255 mOsm, pH 7.4) and fixed in a chamber. 10-15 μL GUV suspension was pipetted onto the surface and incubated for 30 min. Excess lipid material was removed by carefully rinsing with buffer B.

### Indirect FRAP experiments

Indirect FRAP experiments were performed to determine the diffusion coefficient *D*_ΔN49_ of the fusion active acceptor complex ΔN49 in the solid supported part of pore-spanning membranes (s-PSM). A circular region of interest (ROI, *w* = 2.2 μm) was placed on top of the freestanding part of the PSM (f-PSM) and the fluorescence was bleached within this ROI. Since the observed fluorescence recovery in the ROI depends on the diffusion of protein in the s- and f-PSM, *D*_ΔN49_ was obtained by comparing the acquired data with simulated recovery curves of indirect FRAP experiments using FEM simulations. For detailed information see the Supporting Information (Fig. S3).

### Single vesicle assay on PSMs

Single vesicle experiments were performed using an upright spinning disc confocal setup (spinning disc: Yokogawa CSU-X, Rota Yokogawa KG, Wehr, Germany; camera iXon 897 Ultra, Andor technology, Belfast, United Kingdom, pixel size 222 × 222 nm^2^) equipped with a water immersion objective (LUMFLN 60XW 60×, NA 1.1, Olympus, Hamburg, Germany). SRB was excited at *λ*_ex_ = 561 nm and Atto655 DPPE at *λ*_ex_ = 639 nm. Emission light was separated and aligned on either side of the 512 × 512 pixel^2^ detector using an optosplit (Acal BFi Germany, Dietzenbach, Germany) equipped with a 595/40 and 655lp emission filter. Single vesicle docking and fusion events were recorded with a resolution of 20.83 ms/frame over 6.94 min. Each docked vesicle was tagged manually with a minimum 4 × 4 pixel^2^ ROI. Time-resolved fluorescence intensity readout and data evaluation was achieved semi-automatically using a custom-made MATLAB script.

## RESULTS AND DISCUSSION

While lipid mixing assays report on the merging of the outer and inner leaflets of the fusing membranes, no information is gathered about the formation of fusion pores. To address fusion pore formation in our SNARE-mediated single vesicle fusion assay based on pore-spanning membranes (PSMs) (30), we entrapped a water-soluble fluorescent dye into the vesicles at self-quenching concentrations (Fig. 1A).

**FIGURE 1.**
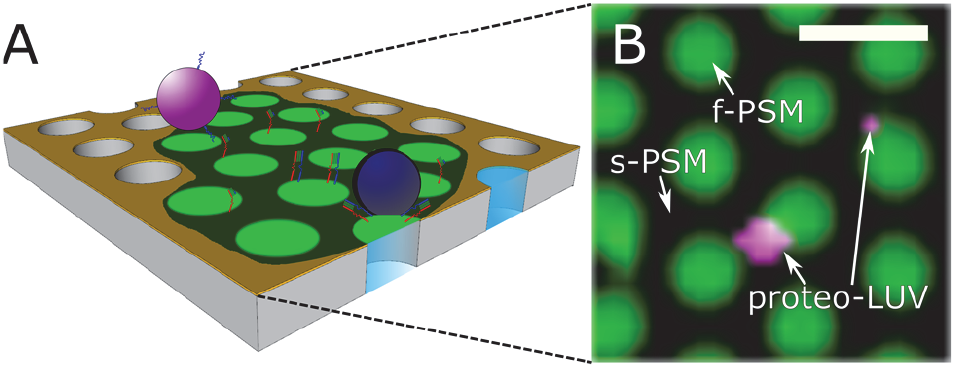
Schematic drawing of the fusion assay based on pore-spanning membranes (PSMs) (A) and fluorescence micrograph (B) of Atto655 DPPE labeled PSMs (false colored in green) containing the t-SNARE complex ΔN49 (DOPC/POPE/POPS/cholesterol (5:2:1:2, *n*/*n*), p:l = 1:500). Proteo-LUVs (false colored in magenta) doped with fulllength syb 2 with the same lipid composition and filled with sulforhodamine B (43 mM) are docked to the PSMs. Scale bar: 2 μm.

To setup the system, we reconstituted the t-SNARE acceptor complex ΔN49 composed of syntaxin 1A (aa 183-288), SNAP25a and the soluble synaptobrevin 2 (syb 2) fragment (aa 49-96) into the PSMs and the v-SNARE synaptobrevin 2 (syb 2) into large unilamellar vesicles (LUVs) (31). PSMs composed of DOPC/POPE/POPS/cholesterol (5:2:1:2, *n*/*n*) and doped with the ΔN49 complex were visualized by fluorescence microscopy using 1 mol% Atto655 DPPE (Fig. 1B).

Owing to the gold covering the top part of the porous substrate, only the freestanding PSMs (f-PSMs) are visible in the fluorescence micrographs, while the supported parts (s-PSMs) appear black as a result of fluorescence quenching (37,38). LUVs composed of DOPC/POPE/POPS/ cholesterol (5:2:1:2, *n*/*n*) doped with syb 2 and filled with SRB were added to the PSMs (Fig. 1B), while taking time series to monitor the docking and fusion of individual vesicles. To measure single vesicle content release (SRB) and lipid mixing (Atto655 DPPE) simultaneously, both fluorescent dyes were recorded with high temporal resolution by means of dual color spinning disc confocal microscopy. SNARE-specificity of docking was proven by blocking the SNARE-binding site of the ΔN49 complex with a soluble syb 2 fragment (aa 1-96) that is known to block fusion prior to protein reconstitution (35). No docking of syb 2 doped LUVs to PSMs was observed under these conditions. Specific docking of syb 2 doped LUVs on the PSMs was observed primarily on the supported parts of the PSMs in agreement with previous observations (29). In contrast to the s-PSMs, which appear black due to the gold induced quenching, the vesicles are sufficiently large so that the fluorophores are far enough away from the gold surface (> 15 nm) to become fluorescently visible (37,38).

### Mobility and reconstitution efficiency of fusion active acceptor complex

The specific docking of syb 2 doped vesicles indicates the successful reconstitution of the ΔN49 complex in the PSMs. However, to quantify the reconstitution efficiency and the mobility of the ΔN49 complex within the PSMs, we performed additional experiments using a ΔN49 complex that was fluorescently visualized with an Atto488-labeled syb 2 fragment (aa 49-96 S79C). In contrast to a study, where SNAP25a was labeled via maleimide chemistry at a single cysteine residue to determine the reconstitution efficiency in GUVs (39), we pursued this approach to ensure that only the fusion active ΔN49 acceptor complex is observed and not the fusion inactive SNAP25a/syntaxin 1A (2:1) complex (35). Fusogenicity of this construct was verified in a bulk fusion assay (Fig. S2B). By using lipid probes as standards (Fig. S2A), the fluorescence intensities of ΔN49-Atto488 doped GUVs (*N* = 1015) were read out to calculate the reconstitution efficiency (Fig. 2A) of the ΔN49 complex. By fitting a lognormal function to the histogram, a reconstitution efficiency of 26 ± 24 % (median) was obtained. In few cases, a reconstitution efficiency of > 100 % was calculated. As the reconstitution of the ΔN49 acceptor complex into SUVs was proven to be quantitative (29) and SUVs are the starting material for the preparation of GUVs, the large variation in protein content is attributed to the electroformation process (29,40,41). Each individual GUV forms one PSM patch on the porous substrate, which explains why we find different numbers of docked vesicles on the individual membrane patches. The average number of docked vesicles was determined to be 0.43 ± 0.56 docked vesicles/μm ^2^.

**FIGURE 2.**
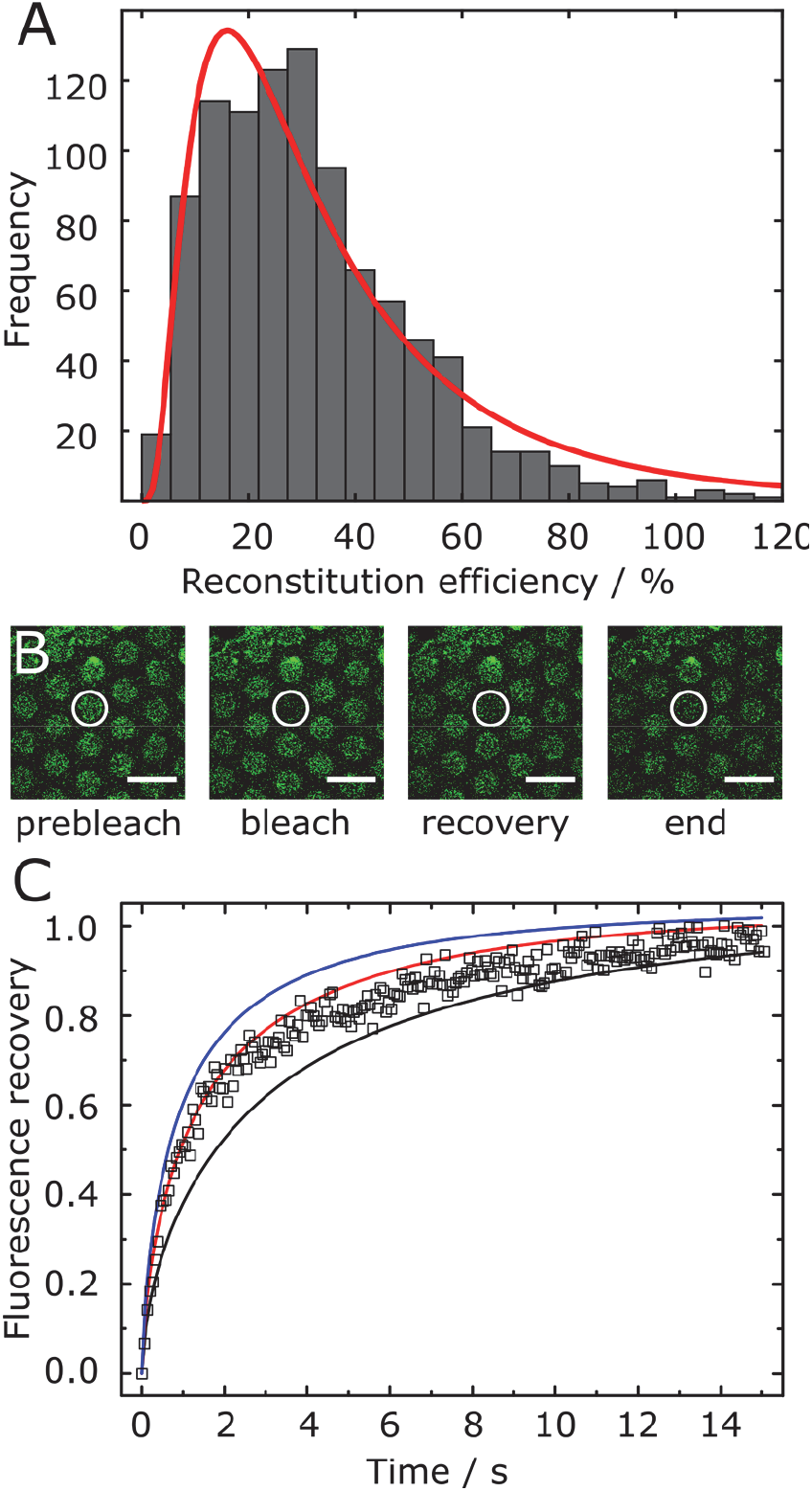
Determination of the reconstitution efficiency and mobility of the ΔN49 acceptor complex labeled with an Atto488 syb 2 (49–96) fragment. (A) Reconstitution efficiency of the ΔN49-complex in giant unilamellar vesicles composed of DOPC/POPE/POPS/cholesterol (5:2:1:2), p/l = 1:500 (*N* = 1015). The red line is a result of fitting a log-normal distribution to the data with a median value of 26 ± 24 %. Values with a reconstitution efficiency > 120 % are not displayed in the graph (*M* = 49). (B) Fluorescence micrographs of an indirect FRAP experiment. The fluorescence of a f-PSM composed of DOPC/POPE/ POPS/cholesterol (5:2:1:2), doped with Atto488 labeled ΔN49 acceptor complex (p:l = 1:250) was bleached (white circle, *r* = 2.2 μm) and the recovery of the fluorescence in the f-PSM was monitored as a function of time. Scale bar: 10 μm. (C) Normalized, mean (*N* = 33), time-dependent fluorescence recovery curve (open squares) and simulated recovery curves with *D*_ΔN49_ (s-PSM) = 0.5 (black), 1 (red), and 1.5 (blue) μm^2^ s^−1^ and *D*_ΔN49_ (f-PSM) = 3.4 μm^2^ s^−1^.

To investigate the lateral mobility of the ΔN49 acceptor complex in the PSMs, we performed so-called indirect fluorescence recovery after photobleaching (FRAP) experiments (Fig. 2B) as described previously (29). By averaging 33 individual FRAP curves (Fig. 2C) and comparing them with simulated recovery curves using finite element simulations (Supporting Information), a diffusion coefficient of *D*_ΔN49_ = 1.0 ± 0.5 μm^2^ s^−1^ and *D*_Lipid_ = 2 ± 1 μm^2^ s^−1^ (Fig. S3B) was estimated. The protein complex mobility is similar to the one that we determined for the single OregonGreen labeled syntaxin 1A transmembrane domain (*D*_syx1A_ = 1.1 ±0.2 μm^2^ s^−1^) (29) and shows that the diffusion coefficient is mainly determined by the transmembrane helix embedded into the bilayer.

### Kinetics of single vesicle content release events

Even though single vesicle fusion assays are rich in information, as each fusion event can be analyzed individually in a time resolved manner, the drawback is that a sufficiently large number of events needs to be evaluated to provide good statistics. Here, we detected 1607 docked vesicles of 7 different preparations and 68 individual membrane patches by their SRB fluorescence. 840 of the 1607 vesicles (52 %) proceeded to fusion with the PSMs. Fluorescence intensity time traces of each docked vesicle within a ROI (Fig. 3A) was read out from the time series. Complete content release upon fusion pore formation is detected by a decrease in SRB fluorescence intensity to the baseline level (Fig. 3A/B). Lack of an increase in PSM fluorescence intensity (Fig. 3B, green) indicates no detectable influx of lipids from the PSM via a fusion stalk. This observation implies a very rapid formation of a fusion pore concomitant with the release of the content dye and collapse of the vesicle into the PSM.

**FIGURE 3.**
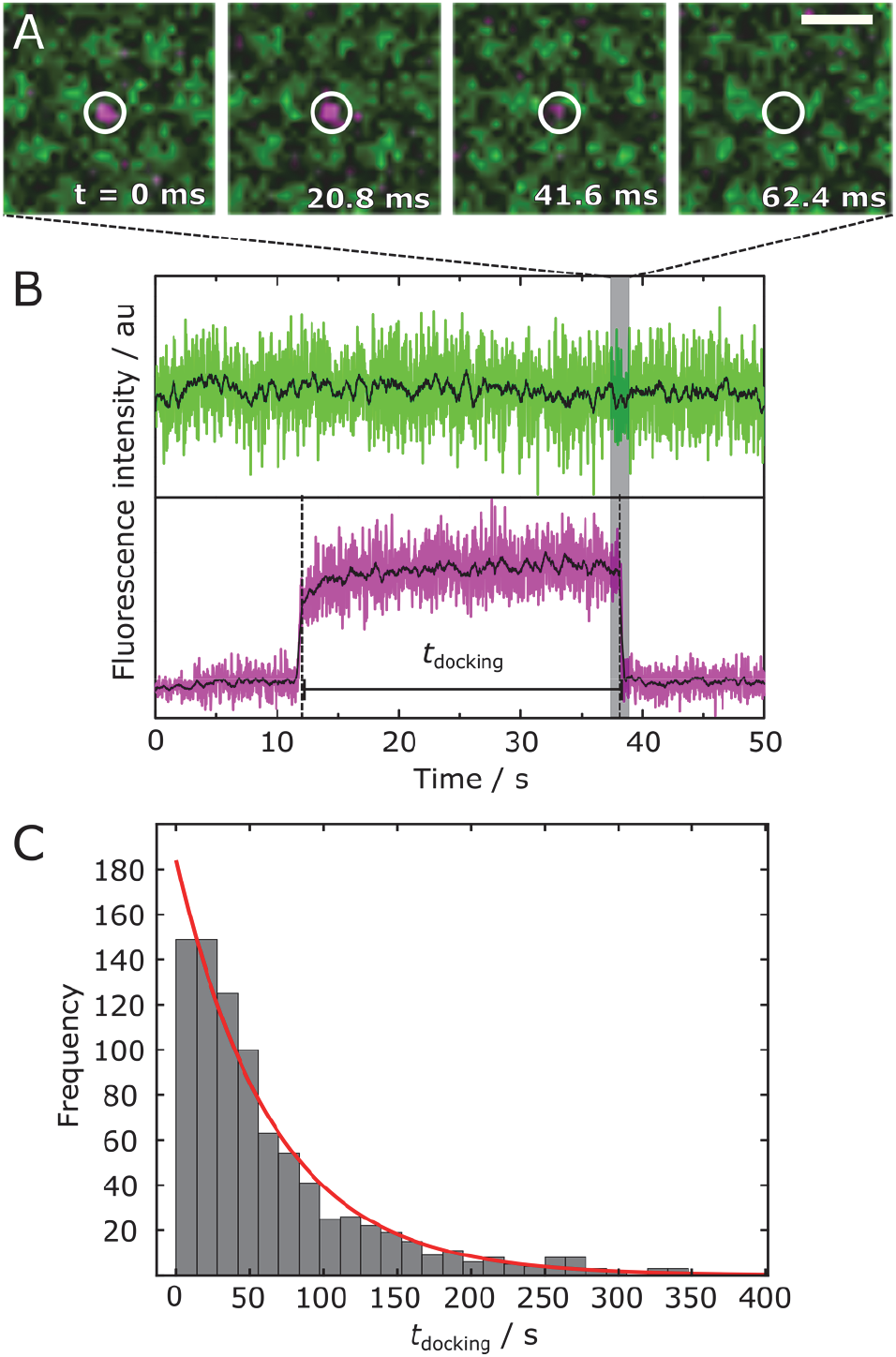
(A) Fluorescence micrographs of a docked vesicle filled with SRB (magenta). The region of interest (ROI) used to readout the fluorescence time is highlighted with a white circle. The vesicle fuses with the PSM (green). Scale bar: 2 μm. (B) Time-resolved fluorescence intensity traces of the vesicle (magenta) and the membrane (green) fluorescence intensity obtained from the ROI in (A). The black lines are 20 data points averaged data. (C) Docking time distribution obtained by extracting *t*_docking_ from *N* = 840 fusion events. A mono-exponential decay function (red) was fit to the data with an average lifetime *τ*_docking_ = 65 ± 4 s.

The majority of fusion assays on the single vesicle level rely on lipid mixing to observe merging of the two membranes (16,29,31,32,42,43). Assays that make use of the release of an entrapped dye in the vesicle lumen to gather information about fusion pore formation are mainly based on planar membranes supported on glass surfaces lacking a second aqueous compartment (20,22,21,44). As concluded by Wang et al. (21), lack of a second aqueous compartment, however frequently leads to vesicle rupturing instead of actual SNARE-mediated content release.

Owing to the porous mesh underneath the PSMs providing an aqueous compartment, in our setup we are able to distinguish between vesicle rupture and fusion pore formation. Upon fusion pore formation the vesicle content becomes visible inside the aqueous compartment underneath the neighboring f-PSM (Fig. S4). In the very rare case of vesicle rupturing (0.003 %, Fig. 6) the vesicle content is, however released into the bulk solution above the membrane (Fig. S5). Such burst is visible as a spike in the SRB fluorescence intensity time trace (Fig. S5B) and can be readily used to separate all burst events from fusion pore formation events. All fusion events were taken to determine the docking time distribution (*N* = 840). Fig. 3C shows the obtained histogram. Fitting a mono-exponential decay function to the histogram results in an average docking lifetime of *τ*_docking_ = 65 ± 4 s. This mono-exponential decay of *t*_docking_, indicating a one step process for fusion, was also observed by Gong et al., who monitored content mixing between two vesicle populations on a single vesicle level (45). The observed docking lifetimes of tens of seconds are similar to those reported previously for synthetic vesicles (29,31) and chromaffin granules (32) on PSMs with reconstituted ΔN49 complex using lipid mixing as a readout parameter.

During the fusion process, several different fusion pathways could be distinguished each being unique in their fluorescence intensity time traces (Fig. 4). While the fluorescence intensity time trace of SRB reports on the formation of a fusion pore releasing the dye into the aqueous compart-ment underneath the PSM, the Atto655 DPPE fluorescence provides information about the influx of this lipid dye into the 3D structure of the fusing vesicle.

**FIGURE 4.**
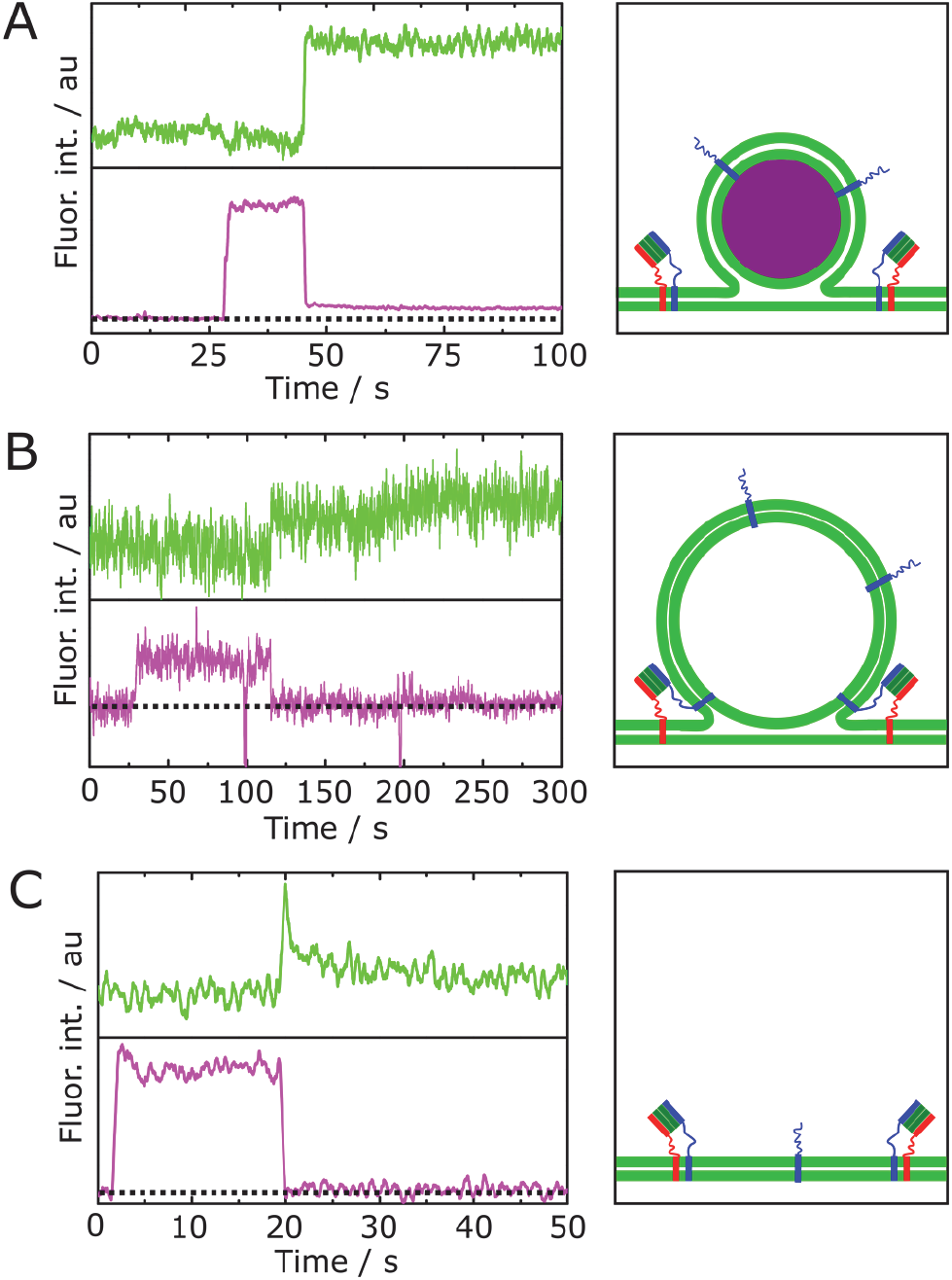
(Left) Fluorescence intensity time traces of fusing vesicles (magenta) with the planar membrane (green) showing different fusion pathways with (right) the corresponding schematics of the (post)fusion structure. (A) Incomplete content release with a remaining three-dimensional (3D) postfusion structure. (B) Complete content release with a remaining stable 3D postfusion structure. (C) Complete content release with an unstable 3D postfusion structure.

An increase in Atto655 DPPE fluorescence intensity indicates that lipid material diffuses into the vesicular membrane thus increasing the distance of the lipid dye to the gold surface, which results in a dequenching of the fluorescence. Only if the lipids are more than 15 nm away from the surface, dequenching is observed indicating that a 3D postfusion structure is present on the PSM surface. In case of an incomplete content release (12 % of all fusion events, see Fig. 6), due to the formation of a metastable fusion pore, the 3D structure remains intact during the whole observation time (Fig. 4A). Besides partial release of the content, also complete release events were observed. Here, we distinguish between two cases: Either a complete release is observed, while the 3D postfusion structure remains intact (Fig. 4B) or the 3D structure becomes instable and fuses into the PSM (Fig. 4C). The cases of a remaining 3D vesicular structure, both full and partial release, is in agreement with other studies described as “kiss and run” fusion (3,46–50). This appears to be one of the possible fusion pathways even in this very simplified fusion system. The parameters that control the different pathways remain, however as yet unclear.

As we are able to determine the time points, where the fusion pore opens and the lipids from the PSM diffuse into the fusing vesicle, we looked more deeply into this kinetics. On the one hand, we determined the time span *τ*_3D_ between Atto655 DPPE influx into the fusing vesicle and the onset of the collapse of the 3D vesicular structure (Fig. 5A/B). Fig. 5C shows the cumulative distribution function of *τ*_3D_ (*N* = 234). Only a bi-exponential functional described the data well with rate constants of *k*_1_ = 0.043 ± 0.004 s^−1^ and *k*_2_ = 0.43 ± 0.01 s^−1^. This result suggests that there are two vesicle populations, one that starts to collapse fast and the other one remaining intact longer. Previously, we reported on such two vesicle populations in a qualitative manner by making use of a lipid mixing assay (29). More importantly, even in live chromaffin cells, two vesicle populations have been reported (51). The kinetic constant obtained for the fast collapsing synthetic vesicles is in good agreement with the kinetics we found for chromaffin granules (32) and is in the same range as values found for the endocytosis kinetics observed *in vivo* (52,53) implying that these type of collapsing vesicles can be directly retrieved from the target membrane in a “kiss and run” manner. Owing to the very high time resolution of 22 ms, we were able to determine Δ*t* being defined as the lag time between the Atto655 DPPE diffusion into the 3D structure and fusion pore formation (Fig. 5A). Fig. 5D shows the corresponding histogram (*N* = 330) of Δ*t*. A positive value of Δ*t* means that the influx of Atto655 DPPE into the 3D vesicular structure occurs prior fusion pore formation. The distribution clearly indicates that in most of the fusion processes content release follows lipid mixing with a median time difference of only 42 ± 11 ms, which is close to the resolution limit of the setup of about 20 ms. It becomes clear that both processes happen almost simultaneously in agreement with previous reports (20,33).

**FIGURE 5.**
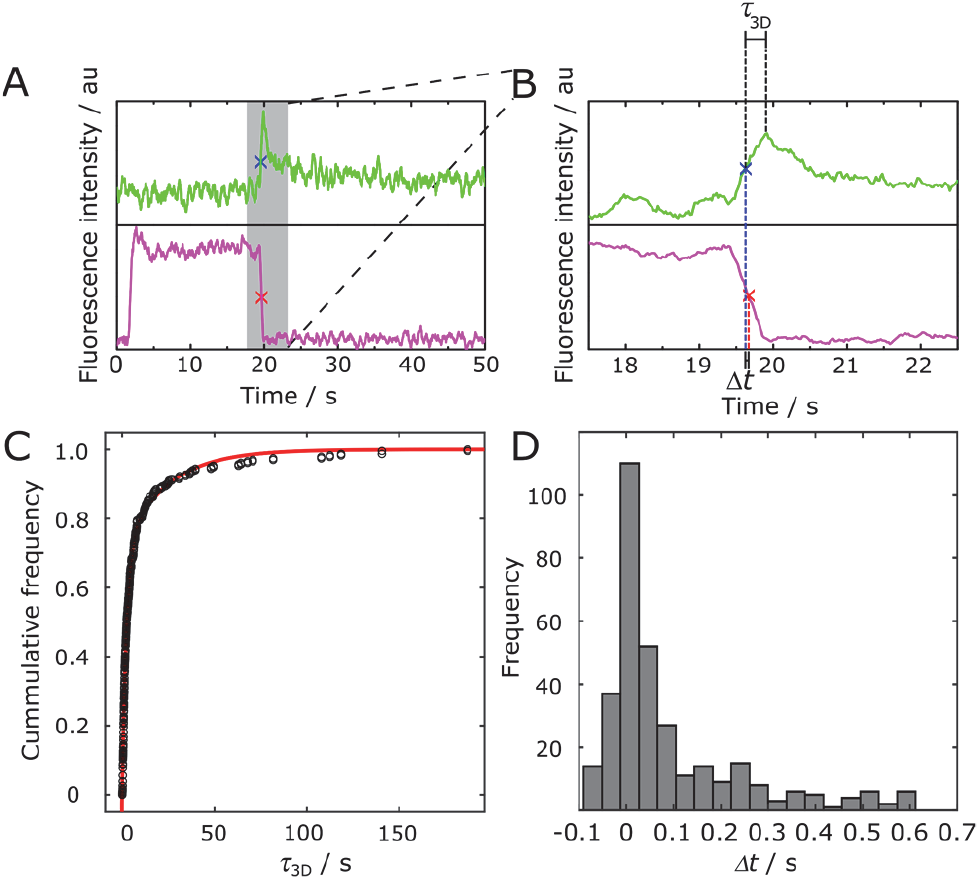
Overview (A) and zoom (B) of time resolved fluorescence intensity traces (smoothed) of a vesicle (magenta) fusing with the s-PSM (green). An increase in green fluorescence intensity is attributed to the Atto655 DPPE diffusion into the 3D structure of the vesicle. *τ*_3D_ is defined as the time span between the Atto655 DPPE influx into the fusing vesicle (marked by a blue x) until the onset of the vesicle collapse. Δ*t* is defined as the lag time between the Atto655 DPPE diffusion into the 3D structure and fusion pore formation (marked by a red x). (C) Cumulative frequency function of *τ*_3D_ (*N* = 234) together with a bi-exponential fit (red) resulting in rate constants of *k*_1_ = 0.043 ± 0.004 s^−1^ and *k*_2_ = 0.43 ± 0.01 s^−1^. (D) Histogram of lag times (Δ*t*, *N* = 330).

In Fig. 6 we summarize all different fusion pathways that we observed. The different pathways were identified according to the two fluorescence intensity time traces (Atto655 DPPE, green and SRB, magenta) (see e.g. Fig. 3B and Fig. 4A-D). Overall, 1607 docked vesicles were analyzed, 52 % of these docked vesicles proceeded to fusion. 24 % of docked vesicles released their content without visible lipid diffusion over a fusion stalk. 452 (54 % of fusing) vesicles underwent fusion with visible lipid mixing into the either stable or unstable 3 D postfusion structure and/or showing an incomplete content release.

**FIGURE 6.**
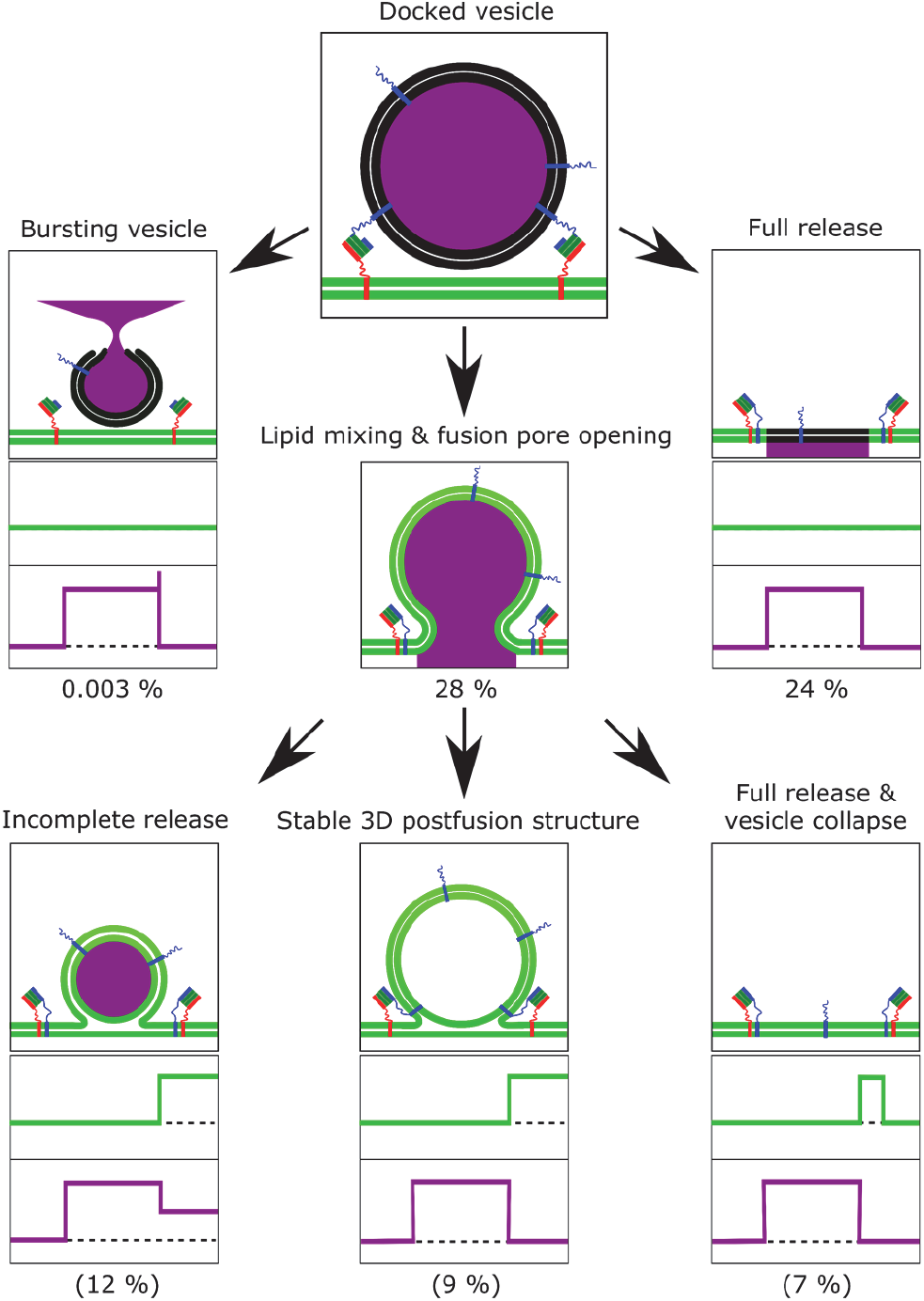
Proposed fusion pathways determined by single vesicle docking/fusion events. 1607 docked vesicles were analyzed, 52 % of these docked vesicles proceeded to fusion. 77 % of these vesicles released their content completely through fusion pore formation. Only 0.003 % of vesicles ruptured and released their content in a burst into the solution above the membrane. Overall 28 % of vesicles fuse via a fusion stalk with different 3D postfusion structures.

77 % of the vesicles that proceeded to fusion released their content completely either directly or in consecutive fusion events. This finding differs from results reported by others, who worked with supported lipid bilayers. Stratton et al. found a large amount of incomplete content release (80 %) (20), while Kreutzberger et al. reported on dead end hemifusion (~ 65%) (22). The results demonstrate that the second aqueous compartment underneath the target membrane alters the ratio of different fusion pathways, presumably reporting on a more physiological scenario with a negligible amount of dead end hemifusion (33,45).

### Flickering fusion pores

Interestingly, in about 50 % of the fusing vesicles, where a first partial content release was observed, the fusion pore re-opens (Fig. 7A/B). In rare cases, this process even consists of three or more consecutive partial dye release events suggesting a “flickering pore”. A flickering pore behavior has been frequently described in *in vivo* experiments (6,54–57). Staal et al. detected consecutive dopamine release events (2-5 times) in midbrain neurons that were proven to originate from one individual vesicle (6). Such fusion pore flickering is thought to first facilitate vesicle recycling in a kiss-and-run manner and would lead to a long-range signaling as well as long-lasting activation of postsynaptic receptors (3,6). In *in vitro* assays, flickering pores have been only rarely described (45). Gong et al. were first to show that stepwise content release is possible in a simple, well defined model system (45). In their study two highly curved vesicle populations containing either the *v*-or the t-SNAREs fused with each other with 72 % fusing in a single step and the remaining 38% fusing in a two- or three-step process. Our findings indeed support the occurrence of multistep release events, which we term flickering pores. Such flickering pores are hence not a function of the full neuronal protein machinery, but can already be observed with the simple minimal fusion proteins reconstituted in a model membrane system. It is, however, likely that the second aqueous compartment facilitates the stepwise fusion pore opening and closing.

**FIGURE 7.**
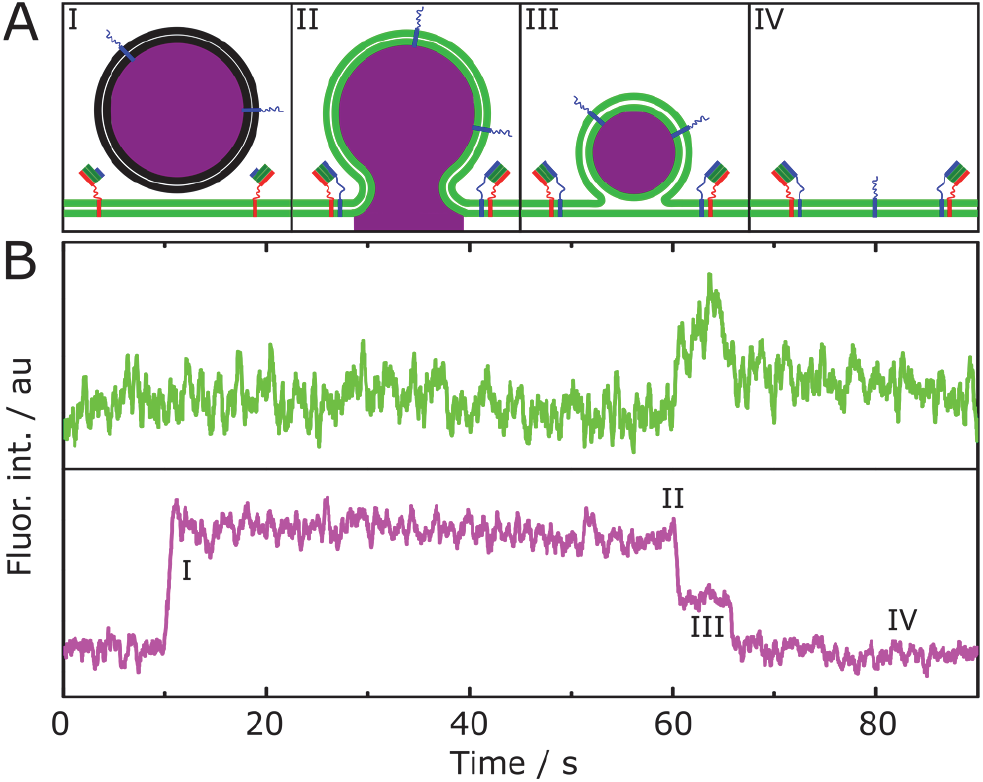
(A) Schematic drawing of the fusion pathway for a stepwise release of the vesicle content with (B) a typical fluorescence intensity time trace of the vesicle (magenta) and planar target membrane (green). Vesicles with a first partial content release (II) undergo a consecutive fusion pore formation with a likeliness of 51 %. 8 % of the whole vesicle population show this flickering pore with 2 % resulting in a complete release as the final situation. In rare cases this flickering pore behavior shows even more than two release events.

## CONCLUSIONS

While several single fusion assays have illuminated elementary steps occurring during neuronal fusion, there is still a lack of *in vitro* assays that are capable of simultaneously reporting on all the different intermediate states including 3D postfusion structures and fusion pore formation including flickering pores without the underlying artifact of vesicle bursting. Owing to the second aqueous compartments underneath pore-spanning membranes, vesicle bursting can be almost completely neglected and can be unambiguously distinguished from fusing vesicles forming a fusion pore. Our results demonstrate that the different fusion intermediates and fusion pathways such as “kiss and run” fusion as well as flickering fusion pores as reported *in vivo*, are apparently not a result of the large number of additional proteins involved in the fusion process, but are already an intrinsic characteristic of the simple neuronal fusion machinery composed of only three SNAREs.

## AUTHOR CONTRIBUTIONS

P.M. performed the fusion experiments. K.H. and L.V. performed the experiments with labeled ΔN49-complex. P.M. and I.M. analyzed the data. C.S. designed the experiments and wrote the manuscript.

## ACKNOWLEDGMENTS

The authors thank the Deutsche Forschungsgemeinschaft for financial support (SFB 803, project B04) and R. Jahn for providing the SNARE constructs.

## SUPPORTING CITATIONS

References (58,59) appear in the Supporting Material.

